# Particle deformability enables control of interactions between membrane-anchored nanoparticles

**DOI:** 10.1101/2023.06.22.546201

**Authors:** Nikhil Nambiar, Zachary A. Loyd, Steven M. Abel

## Abstract

Nanoparticles adsorbed on a membrane can induce deformations of the membrane that give rise to effective interactions between the particles. Previous studies have focused primarily on rigid nanoparticles with fixed shapes. However, DNA origami technology has enabled the creation of deformable nanostructures with controllable shapes and mechanical properties, presenting new opportunities to modulate interactions between particles adsorbed on deformable surfaces. Here we use coarse-grained molecular dynamics simulations to investigate deformable, hinge-like nanostructures anchored to lipid membranes via cholesterol anchors. We characterize deformations of the particles and membrane as a function of the hinge stiffness. Flexible particles adopt open configurations to conform to a flat membrane, whereas stiffer particles induce deformations of the membrane. We further show that particles spontaneously aggregate and that cooperative effects lead to changes in their shape when they are close together. Using umbrella sampling methods, we quantify the effective interaction between two particles and show that stiffer hinge-like particles experience stronger and longer-ranged attraction. Our results demonstrate that interactions between de-formable, membrane-anchored nanoparticles can be controlled by modifying mechanical properties of the particles, suggesting new ways to modulate the self-assembly of particles on deformable surfaces.

## 1 Introduction

The adsorption of nanoparticles onto a lipid bilayer can cause local deformations of the membrane. The local curvature induced by the particle causes an increase in the bending energy of the membrane, and the deformation field of one particle can interact with deformations caused by other nanoparticles in its vicinity. ^1^ These membrane-mediated interactions can lead to effective interactions between the nanoparticles. In some cases, the interaction of membrane deformations can lead to attractive interactions between particles. This can promote their self assembly, ^2^ and in some cases, lead to large-scale deformations of the membrane.^3–6^ Physically, the attraction between particles arises largely due to the total bending energy of the membrane being lower when the particles aggregate.

Studies of membrane-mediated interactions between rigid nanoparticles have characterized how the membrane bending energy depends on the distance between particles, thus leading to a membrane-mediated force between particles that changes as a function of distance.^7–10^ For example, Reynwar et al. showed that spherical colloids experience an attractive force when sufficiently close due to changes in the bending energy of the membrane. ^7^ Membrane-mediated interactions can facilitate self-assembly of particles,^11^ and it has been shown that spherical particles adsorbed to planar and vesicular membranes can form linear aggregates and induce tubulation of the membrane.^4,5,8^ For anisotropic particles, membrane-mediated interactions depend on the relative orientation of the particles,^3,12,13^ and changing the particle shape or properties of the membrane can lead to assembly in different configurations.^6,14^ Curved particles have been of particular interest, in part due to their similarity to BAR proteins, and can induce curvature that leads to mutual interactions, self-assembly, and membrane sculpting.^14–16^

DNA origami nanoparticles have emerged as a valuable tool to study the interactions of membranes and nanoparticles.^17^ DNA origami can be designed with a wide range of well-defined shapes, and they can be readily modified to incorporate cholesterol tags that anchor the particles to membranes. Rigid DNA origami structures assemble on membranes,^18,19^ and curved DNA origami structures have been shown to induce morphological changes of membranes.^20,21^ DNA origami with cholesterol anchors can also produce synthetic pores in lipid membranes^22–24^ and act as nanoscale sensors to measure membrane surface charges ^25^ and transmembrane potentials.^26^

An exciting recent development is the design of deformable DNA origami nanostructures with controllable mechanical features.^27^ DNA origami hinges are an example in which the stiffness of the hinge can be controlled by small changes in the design.^27,28^ Moreover, DNA origami hinges have been designed to be actuated by temperature, ions, and magnetic fields.^29–31^ They also have applications as nanoscale devices, as demonstrated recently with cholesterol-anchored, flexible DNA hinges that act as nanoscale curvature sensors.^32^

Deformable DNA origami nanoparticles open new avenues for studying particle-membrane interactions and resulting membrane-mediated interactions between particles. This is because the particles and membrane can mutually deform one another, introducing new complexities into the problem. However, there have been few efforts to date that illuminate the impact of particle deformability. Previous studies exploring interactions between semiflexible polymers and membranes revealed a complex interplay between deformations of the polymer and membrane.^33,34^ We recently used continuum models and Monte Carlo simulations to study hinge-like particles adhered to an elastic membrane via a short-ranged attractive potential. We showed that strong adsorption promotes attractive, membrane-mediated interactions between particles, and that configurations of aggregates are influenced by hinge stiffness.^35^ However, interactions of deformable, cholesterol-anchored particles in realistic membrane systems have not yet been explored.

In this paper, we use coarse-grained molecular dynamics simulations to investigate hinge-like nanoparticles that interact with a realistic lipid bilayer by means of cholesterol anchors. The purpose of the study is to capture the key physical characteristics of a hinge-like particle and systematically understand the impact of the stiffness of the hinge on particle-membrane and particle-particle interactions. We first briefly introduce the model and investigate isolated particles, both in water and in contact with a lipid bilayer. We then characterize two particles on a membrane using unbiased simulations before using umbrella sampling to quantify the effective interaction between two particles. We conclude by discussing the cooperative behavior observed when particles aggregate and how a new degree of freedom — the mechanical properties of a deformable particle — can be used to control membrane-mediated interactions between particles. Simulation details can be found at the end.

## 2 Results and discussion

We utilized molecular dynamics simulations to probe the behavior of deformable, hinge-like nanoparticles anchored on a lipid bilayer. We employed a coarse-grained model, which allowed us to probe the length and time scales needed to assess particle-induced deformations of membranes and interactions between particles. Specifically, we used the Martini 2.2 coarsegrained force field^36^ because of its success in representing lipids and other biomolecules. Standard Martini representations were used for the phospholipids comprising the lipid bilayer and cholesterol anchors on the hinge-like particle. We created a model of a hinge-like particle that captures key physical attributes of a deformable nanoparticle with realistic cholesterol anchors (Figure 1). It was not intended to represent a real nanoparticle, but to provide general insight into the behavior of a class of deformable, hinge-like nanoparticles. Each of the three hinge angles was subject to a cosine angular potential characterized by a force constant *k* that we varied from 0 to 1000 kJ/mol. The natural angle of the hinge (*θ*_0_) was 120°. Details of the simulations can be found in Methods.

**Figure 1.**
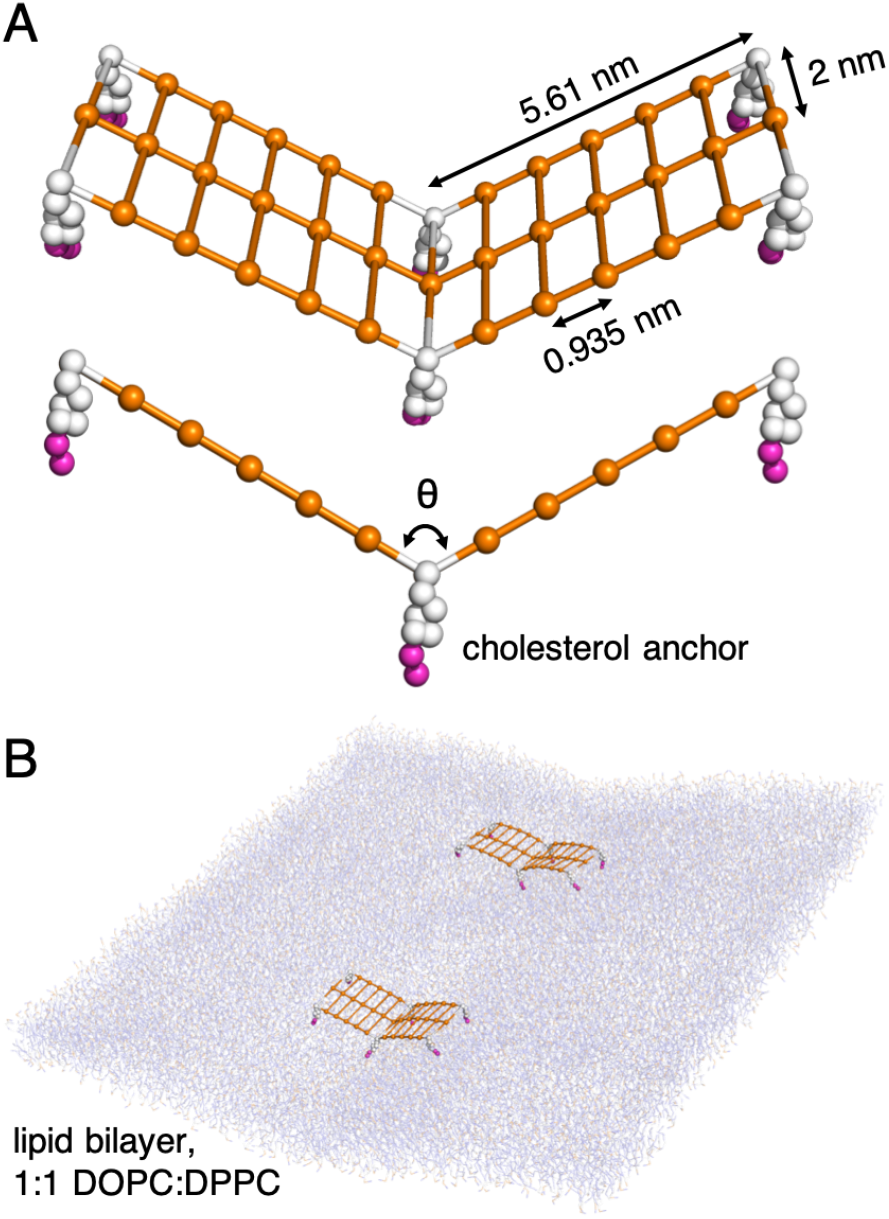
Schematic of the model. (A) Two views of the coarse-grained nanoparticle. Two “arms” are connected in a hinge-like configuration with a natural angle of 120°. The stiffness of the hinge is systematically varied. Six cholesterol anchors (white and fuchsia) serve to anchor the particle to a lipid bilayer. (B) Snapshot showing two particles anchored to a lipid bilayer.

### 2.1 Isolated hinge-like particles

We first sought to characterize the deformable, hinge-like particle in water without a lipid bilayer present. To this end, we ran unbiased simulations to evaluate equilibrium behavior at different values of the hinge stiffness (*k*). Figure 2 shows the distributions of angles sampled by the hinge. At large values of *k*, the distributions are centered at the natural angle of 120°. The width of the distribution increases as the stiffness decreases, indicating that the hinge samples a broader distribution of angles when the angular potential is weaker, as expected.

**Figure 2.**
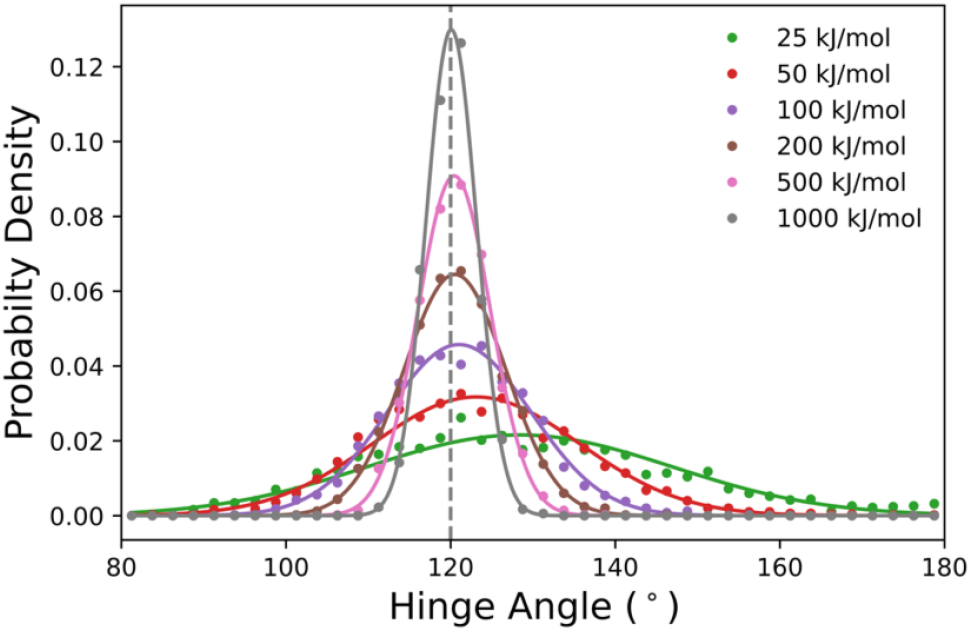
The probability density of the hinge angle (*θ*) at equilibrium for a hinge in water (no bilayer). Different values of the hinge stiffness (*k*) are shown. The vertical dashed line corresponds to the natural angle of the hinge (*θ*_0_). Solid lines are Gaussian fits to the data (circles) meant to guide the eye.

For smaller values of *k* (≲ 25 kJ/mol), the distribution of angles shifts toward larger angles. This corresponds to more open configurations of the hinge. The effect is modest for *k* = 25 kJ/mol (Figure 2), and the effect is most pronounced when *k* = 0 (Figure S1). For this case, the maximum of the distribution is at 180°, which is likely due to electrostatic repulsion between charged beads constituting the arms of the hinge. The open configuration reduces unfavorable interactions, but the effect is sufficiently weak that only the most flexible hinges are substantively impacted. We also assessed whether the presence of cholesterol anchors impacted the distribution of hinge angles by conducting simulations in which the cholesterol anchors were removed. The removal of the anchors resulted in only small changes to the angles sampled by the hinge (Figure S2), indicating that they did not significantly perturb the behavior of the particle in water.

### 2.2 Stiff hinges deform membranes

To characterize the behavior of membrane-anchored particles, we first studied an isolated particle with its cholesterol anchors inserted in a lipid bilayer. We used unbiased simulations to assess how the membrane impacts configurations adopted by the hinge. Figure 3 shows sample configurations for hinges with different values of the bending stiffness. At a small value (*k* = 25 kJ/mol), the hinge appears nearly flat as it conforms to the flat lipid bilayer. For larger values, the hinge no longer appears flat, and it adopts configurations with angles closer to the natural angle of the hinge (120°). This coincides with a large-scale deformation of the lipid bilayer in the region near the hinge. The hinge is closest to its natural angle for the largest value of *k*.

**Figure 3.**
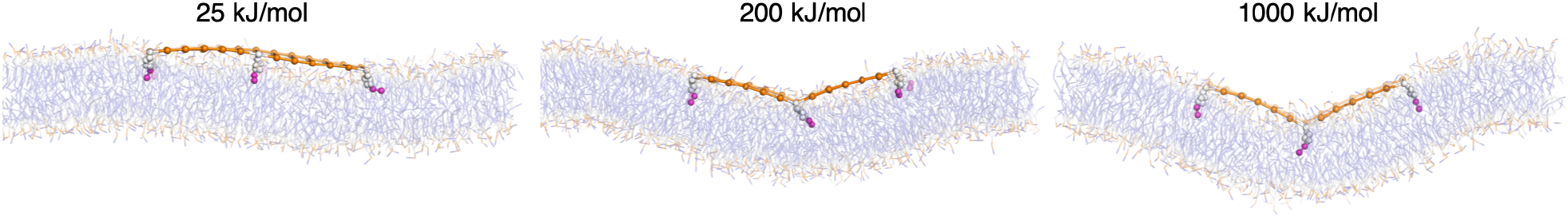
Typical equilibrium snapshots of a hinge-like particle anchored to a membrane. Three values of the hinge stiffness are shown (*k* = 25, 200, and 1000 kJ/mol).

Figure 4 shows the influence of the membrane on configurations of the hinge using the distribution of angles sampled by a membrane-anchored particle. Weaker hinges (*k* ≤ 100 kJ/mol) are significantly deformed by the membrane, with an average angle of approximately 170°. In this regime, the hinge is flexible enough that it is energetically favorable for it to adopt a significantly deformed flat configuration while maintaining a relatively flat membrane shape. In contrast, stiffer hinges (*k* ≥ 200 kJ/mol) adopt configurations that are closer to the natural angle of the hinge. The distribution of angles for the stiffest hinges (*k* = 500 and 1000 kJ/mol) are only slightly shifted compared with the distributions in water, while the intermediate stiffness (*k* = 200 kJ/mol) is intermediate between the natural angle and the fully open hinge. In this regime, the membrane conforms to the particle shape, resulting in particle-scale deformations of the membrane. Thus, membrane deformations can be controlled by the stiffness of the hinge.

**Figure 4.**
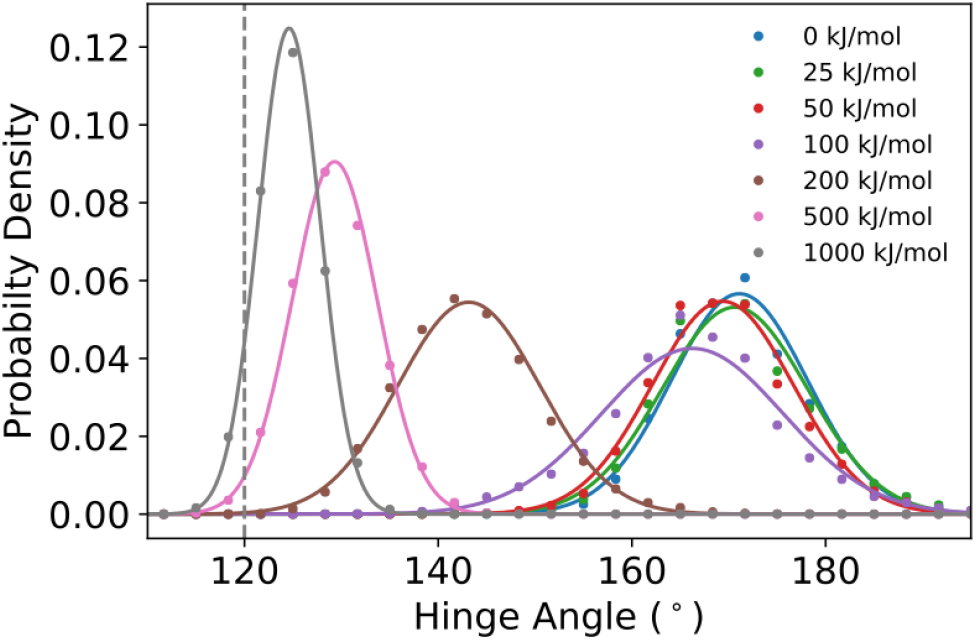
The probability density of the hinge angle (*θ*) at equilibrium for a hinge anchored to a lipid bilayer. Different values of the hinge stiffness (*k*) are shown. The vertical dashed line corresponds to the natural angle of the hinge (*θ*_0_). Solid lines are Gaussian fits to the data (circles) meant to guide the eye.

### 2.3 Membrane-anchored hinges spontaneously aggregate

Studies of rigid nanoparticles have shown that membrane deformations induced by their adsorption can lead to effective interactions between the adsorbed particles. The results above demonstrate that the model hinge-like particle can induce deformations of a lipid bilayer, and that the extent of the deformation can be controlled by the stiffness of the hinge. Thus, we expect the hinges to experience deformation-mediated interactions. However, in contrast with rigid particles, the presence of more than one hinge can lead to cooperative effects in which hinges mutually influence one another’s configurations.

To investigate membrane-mediated interactions between particles, we first conducted unbiased simulations of two hinges anchored to the lipid bilayer. Figure 5 shows the distance between the centers of mass of the two hinges as a function of time for various values of the hinge stiffness. We considered two sets of initial configurations: one in which the hinges started far apart, and one in which they started close together. When starting far apart (Figure 5A), the distance between particles fluctuates but, for most cases, eventually exhibits a rapid decrease to a stable plateau of ≈ 5 nm. This distance is consistent with them being side-by-side in close proximity. When starting close together (Figure 5B), the hinges stay in close proximity for all hinge stiffnesses, again indicating an effective attraction. These results suggest that, when sufficiently close, the two particles experience an effective attraction that causes them to aggregate, and that the range of the attraction is larger for larger values of *k*.

**Figure 5.**
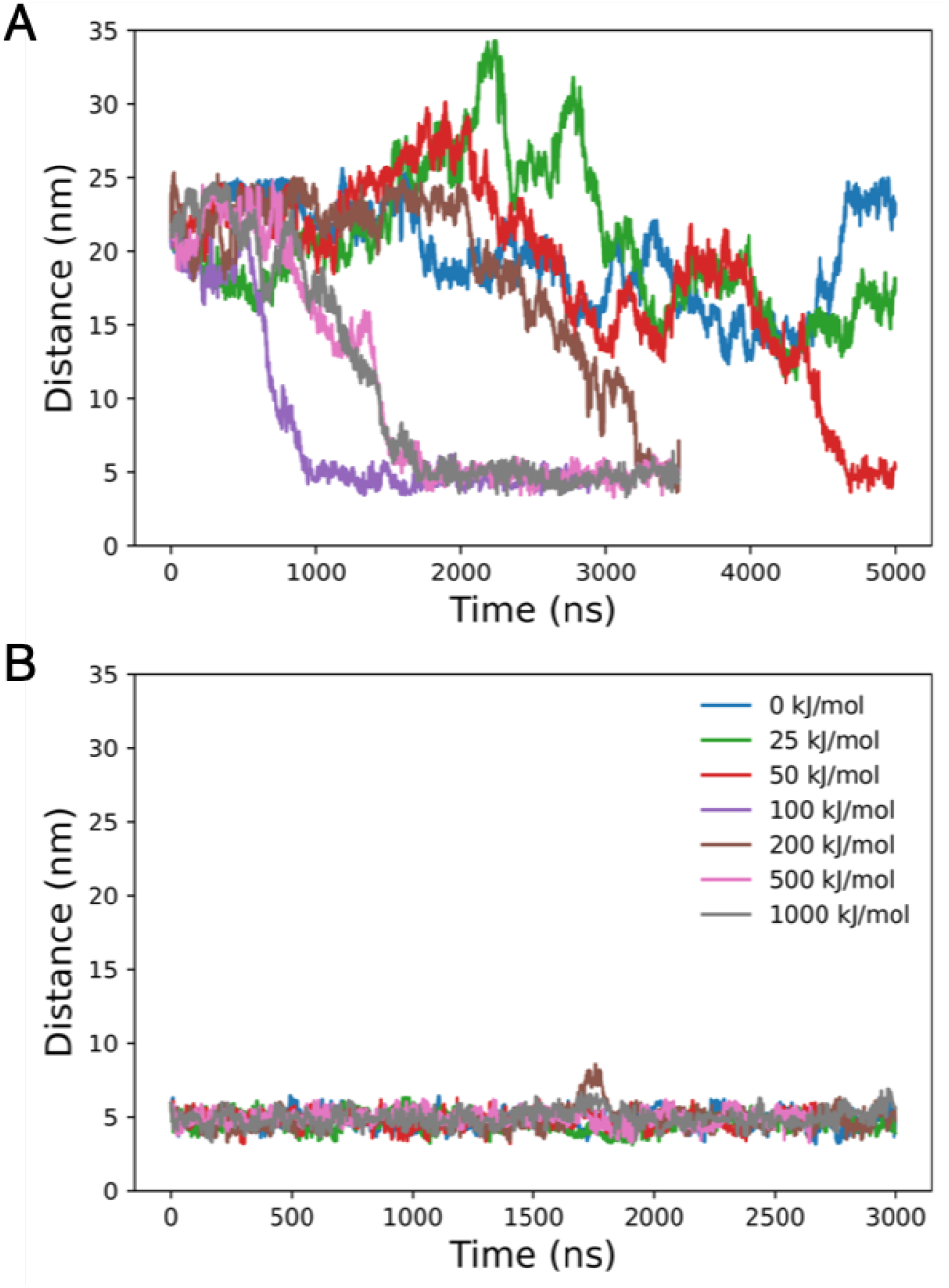
Distance between centers of mass of two particles anchored to a lipid bilayer as a function of time. Hinges are started far apart in (A), with the initial distance equal to half the box size. Hinges are started close together, in side-to-side contact, in (B).

Figure 6 shows snapshots taken from two simulation trajectories with weak and strong hinges, respectively. The locations of the hinges are superimposed on a heatmap indicating the deviation of the membrane from its average position in the out-of-plane dimension (Δ*z*).

**Figure 6.**
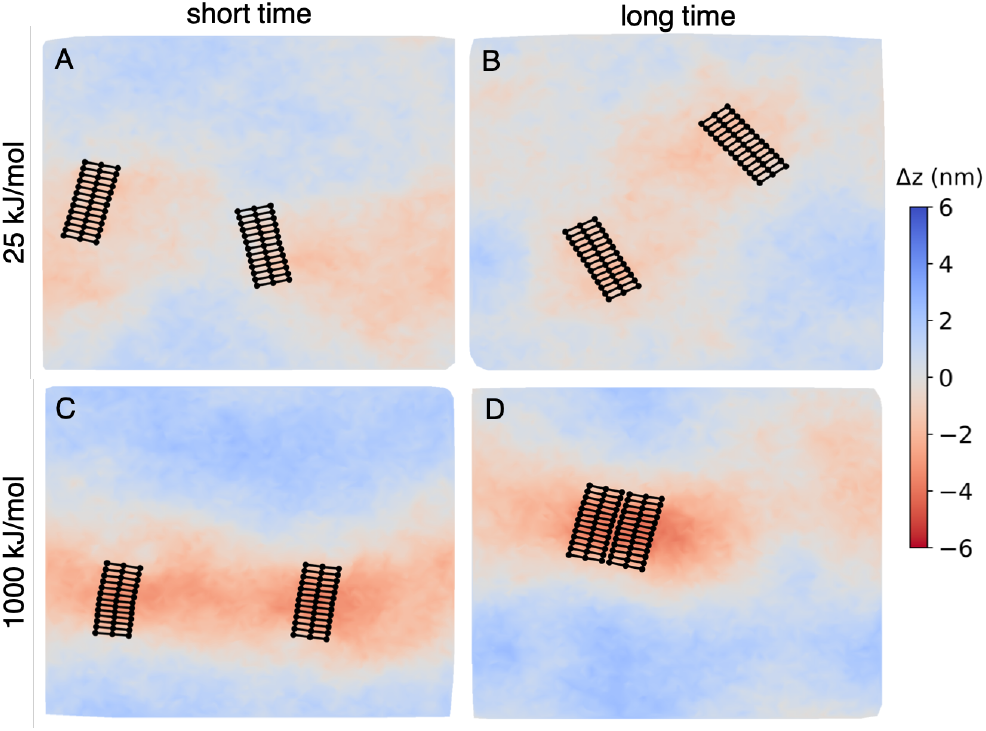
Top-down view of hinges and deviations of the membrane height. Δ*z* denotes the deviation of the membrane from its average position in the out-of-plane dimension. (A),(B) Snapshots of weak hinges (*k* = 25 kJ/mol) at 50 ns and 3 *μ*s, respectively. (C), (D) Snapshots of strong hinges (*k* = 1000 kJ/mol) at 50 ns and 3 *μ*s, respectively.

This demonstrates the deformations induced by the hinges. The stiffer hinges induce larger deformations of the membrane, and the furrow-like nature of the deformation can be seen by the extension of the red color (Δ*z <* 0) orthogonal to the direction of the arms of the hinge. For the stiff hinges, the deformation appears more localized around the particles when they aggregate.

The overall energy of the system is impacted by both the energy of deforming the hinges and the energy of deforming the membrane; the previous results suggest that it is energetically favorable for the two hinges to be in close proximity. To assess cooperative effects when two hinges are present, we analyzed the angles sampled by two membrane-anchored hinges when they started in close proximity (corresponding to Figure 5B). Figure 7 compares the behavior of two aggregated hinges to an isolated hinge anchored to the bilayer. When two hinges are in close proximity, the average angle of the hinge decreases, reflecting a configuration of the hinge that is closer to its natural angle of 120°. The cooperative effects are most pronounced at intermediate values of *k*. When *k* is small, the hinges remain almost completely flat and induce relatively little membrane deformation whether there are one or two hinges. When *k* is large, a hinge is sufficiently stiff to significantly deform the membrane, whether on its own or in conjunction with another hinge. At intermediate values of *k*, two hinges together allow the hinges to share the energetic cost of deforming the membrane, and each adopts a configuration closer to its natural hinge angle. Two hinges together thus cause a larger deformation than when only one isolated hinge is adsorbed to the membrane.

**Figure 7.**
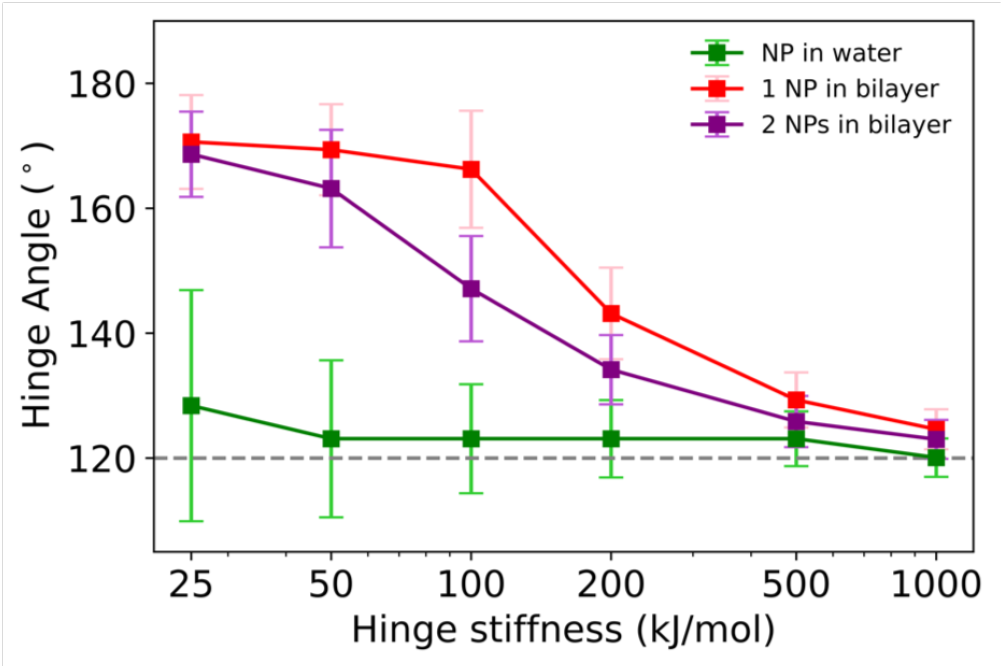
Comparison of the angles sampled by an isolated hinge in water, an isolated hinge anchored to a membrane, and two hinges anchored to a membrane when started in close proximity. The horizontal dashed line corresponds to the natural angle of the hinge (*θ*_0_).

### 2.4 Hinge stiffness controls interactions between membrane-anchored particles

The spontaneous aggregation of two particles and the resulting stability of the aggregate demonstrate an effective attraction between two membrane-associated particles. To characterize the nature and magnitude of the effective interaction, we used umbrella sampling methods to calculate the potential of mean force (PMF) between the centers of mass of the two particles in the plane of the membrane.

Figures 8 and S3 show multiple PMFs, each of which was obtained for two hinges of a particular stiffness. A general feature is a minimum in the PMFs at a distance of ≈ 5 nm. This corresponds to the distance at which the hinges are in side-by-side contact, as demonstrated in Figure 6. Starting at larger distances, there is an effective attraction between the two hinges, and there are no appreciable energy barriers to prevent the spontaneous aggre-gation of the particles. A short-ranged (*<* 5 nm) repulsion is associated with short-ranged repulsive interactions of the beads comprising the hinges. These results confirm that it is energetically favorable for the hinges to aggregate spontaneously.

**Figure 8.**
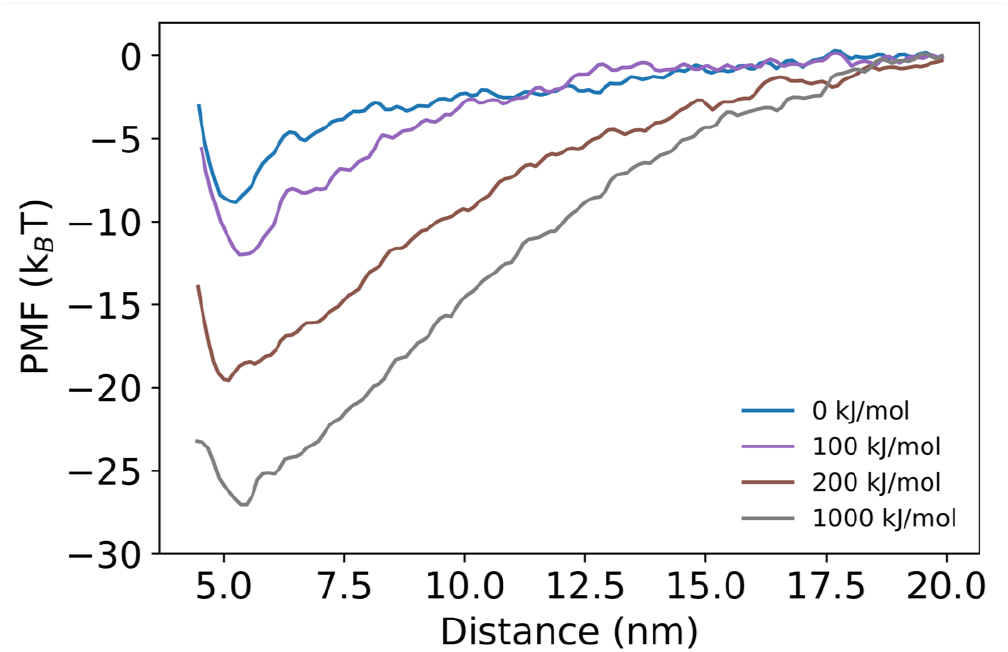
Potential of mean force as a function of the distance between the centers of mass of the two hinge-like particles in the plane of the membrane (*xy*-plane). Additional hinge stiffnesses are omitted for clarity and can be found in Figure S3.

A key feature of Figure 8 is that stiffer hinges result in stronger attraction, as revealed by the increasing depth of the PMF. Further, stiff hinges experience an appreciable attractive force at larger distances than the weaker hinges. This is consistent with the faster aggregation of stiffer particles in unbiased simulations. Physically, stiffer hinges cause larger deformations of the membrane, thus increasing the range of the bending and the total bending energy of the membrane.

The physics underlying the effective interaction between hinges is that the total deformation of the membrane is decreased when two hinges are close together, thus decreasing the bending energy of the membrane. This decrease in bending energy of the membrane is the driving factor for the aggregation of the hinge particles. We note that a modest attractive interaction is present even when the hinge is freely jointed (*k* = 0). This is likely because the particle becomes slightly distorted when the cholesterol anchors insert into the membrane (Figure S4), thus impacting the local shape of the membrane. The results indicate that the overall energy is decreased when the particles are in close proximity, leading to an effective attraction. Our focus in this paper is on the effects of hinge stiffness, which clearly modulates the magnitude of membrane deformations and strength of attraction between particles. This demonstrates that the degree of particle deformability can used to tune membrane-mediated interactions between particles.

## 3 Conclusions

In this work, we studied deformable nanoparticles anchored by cholesterol to a realistic lipid membrane using coarse-grained computer simulations. Motivated by a new class of DNA origami designs,^27,28^ we used a generic model of a hinge-like nanoparticle that allowed us to systematically change the stiffness of the hinge region. We showed that the particle and membrane could mutually deform one another: When the hinge was highly flexible, it adopted an open configuration and conformed to the flat shape of the membrane. As the hinge stiffness increased, it produced progressively larger deformations of the membrane. Mutual deformations were most pronounced at intermediate values of the stiffness, where both the particle and membrane were significantly deformed compared to their typical equilibrium configurations. We then studied membrane-mediated interactions between two particles, showing that two particles would spontaneously aggregate in a side-by-side configuration. When assembled this way, the deformations induced by the particles were aligned and formed a trough-like deformation perpendicular to the arms of the hinge. Aggregated particles also adopted configurations closer to their natural hinge angle and thus collectively induced larger deformations of the membrane. This effect was most pronounced at intermediate hinge stiffnesses.

A key result is shown in Figure 8, which demonstrates that the strength and range of attraction between two particles increases with increasing hinge stiffness. This demonstrates that membrane-mediated forces acting on deformable particles can be controlled by changing mechanical properties of the particles. Thus, membrane-mediated self-assembly of particles can be influenced by mechanical properties of the particles alone. This further suggests that actuating the mechanical properties of the particles, for example by external perturbation,^29–31^ could provide a means to control particle-particle interactions in a time-dependent and externally controlled manner. The results further show cooperative effects when particles aggregate, whereby particles change their configurations because they can collectively deform the membrane. FRET-based measurements may be useful for assessing changes in hinge configurations,^32^ and it will be interesting to study how self-assembly in large-scale systems is impacted by particle deformability and cooperative effects.

## 4 Methods

### 4.1 Model of the hinge-like particle

We created a coarse-grained nanoparticle inspired by DNA origami hinges. The model was not intended to represent an actual DNA origami particle, but to capture key physical characteristics in a manner that allowed us to systematically investigate its behavior when anchored to a membrane. The size of the particle was smaller than existing DNA origami hinge designs, which have longer arms, for computational tractability. The primary effect of size is to change the scale of the membrane deformations but not the underlying physics of the membrane-mediated interactions.

The nanoparticle consisted of two stiff arms connected at an angle to form a hinge as shown in Figure 1. Each of the arms consisted of 3 rows of 6 beads. The beads in each row were spaced 0.935 nm apart. The rows were spaced 2 nm apart, which is consistent with the diameter of a DNA double helix. All beads in the arms were connected in a rectangular lattice via harmonic bonds with force constant 12, 500 kJ mol^*−*1^ nm^*−*2^. Cosine-based angle potentials were also imposed, with all 90° angles having a bending constant of 25,000 kJ/mol and all 180° angles having a bending constant of 250,000 kJ/mol.

Cholesterol anchors were included by incorporating the Martini SP1 bead (representing the portion of cholesterol containing the hydroxyl group) into the arms of the nanoparticle at the ends and center of rows 1 and 3. The rest of the beads in the arm were Martini beads of type Q0, which were used to mimic the phosphate linkages on the outside of a DNA double helix. A cosine-based angle potential was used to set the hinge stiffness: *V*_*a*_(*θ*) = *k* cos(*θ*) −cos(*θ*_0_))^2^*/*2. The natural angle of the hinge (*θ*_0_) was chosen to be 120°, and the force constant for the cosine potential was varied from 0 to 1000 kJ/mol. This potential was used for the angle defined at the middle bead along each of the three rows of beads.

The topology file for the hinge-like particle is included as Supporting Information.

### 4.2 Simulation details

In the MD simulations, we used standard Martini parameters for all non-bonded interactions.^37^ The LINCS algorithm was used for bonded interactions.^38^ Lennard-Jones interactions were cut off at 1.1 nm using the potential-shift-Verlet modifier. Coulombic interactions were also cut off at 1.1 nm using the reaction field method. All systems were solvated with standard Martini water, without utilizing anti-freeze beads. All MD simulations were performed in GROMACS 5.1.2.^39,40^

For simulations of the hinge-like particle in water without a lipid bilayer, we solvated the particle in a 24 nm × 24 nm × 24 nm box. The system consisted of 109,221 beads. First, energy minimization using steepest descent was performed for 5,000 steps. Subsequently, the system was equilibrated in four stages while increasing the simulation timestep from 2 fs to 20 fs. During equilibration, the temperature coupling was handled by the velocity-rescaling thermostat^41^ with a reference temperature of 310.15 K. For all equilibration runs, we used isotropic Berendsen pressure coupling ^42^ with a reference pressure of 1 bar and compressibility of 3 × 10^*−*4^ bar^*−*1^. In production runs, we used the velocity-rescale thermostat and the Parrinello-Rahman barostat.^43^ The production runs were simulated for 2 *µ*s and the final 0.5 *µ*s was used for analysis.

For unbiased simulations with hinge-like particles and a lipid bilayer, we used the CHARMM-GUI Martini maker^44,45^ to first set up a solvated bilayer with a composition of 1:1 DOPC:DPPC. The bilayer size was 25 nm × 25 nm (2,000 lipids) and 50 nm × 50 nm (7,536 lipids) for simulations with one and two hinges, respectively. The dimension orthogonal to the lipid bilayer in both cases was 24 nm. The bilayer was equilibrated using the standard multi-step equilibration procedure from CHARMM-GUI. The equilibration consisted of four stages, where the restraints on the bilayer head groups were relaxed and the simulation time step was increased from 2 fs to 20 fs. One or more hinge particles were then inserted at random locations in the water phase by replacing water beads. The system with one hinge consisted of 132,283 beads; the system with two hinges consisted of 455,019 beads. The standard multi-step equilibration process from CHARMM-GUI was again used for energy minimization and equilibration. MD simulations were run to allow the hinge(s) to diffuse into contact with the lipid bilayer. Once the cholesterol anchors inserted into the lipid bilayer, the system was equilibrated for an additional 2.5 *µ*s.

Trajectories generated from GROMACS were analyzed in python using the MDanalysis package.^46,47^ PyMOL was used to create snapshots of simulations.

### 4.3 Umbrella sampling

We used umbrella sampling methods to determine the potential of mean force (PMF) as a function of the distance between the centers of mass of the two membrane-anchored particles in the *xy*-plane. The final snapshot of the unbiased simulation with hinge stiffness 1000 kJ/mol was used as the initial input. For each hinge stiffness, this snapshot was used as the initial configuration and equilibrated for 1 *µ*s while using a harmonic potential of 1000 kJ mol^*−*1^ nm^*−*2^ centered at the prescribed distance between their centers of mass. The hinges were then pulled away from each other in the plane of the membrane (*xy*-plane) to generate initial snapshots for the umbrella windows. To do this, the natural distance of the harmonic potential was increased at a rate of 0.001 nm/ps. We used a window spacing of 0.45 nm to choose initial configurations and then performed simulations for each window with a force constant of 100 kJ mol^*−*1^ nm^*−*2^ for the harmonic umbrella potential. Each umbrella window was simulated for 1 *µ*s, and the last 0.5 *µ*s was used to calculate the PMFs. To sample configurations when the hinges were close together, we repeated the same procedure by creating snapshots while pushing the particles closer together. Data from each hinge stiffness was analyzed using the weighted histogram analysis method (WHAM)^48^ implemented in GROMACS.^49^ A correction term^50,51^ of *k*_*B*_*T* ln(*R*), where *R* is the distance between centers of mass in the *xy*-plane, was added to the PMF obtained from WHAM to account for the size of state space being a function of *R*.

## Acknowledgement

This work was supported by the National Science Foundation (CBET-2217777).

## Supporting Information Available

Additional simulation results and histograms obtained from umbrella sampling. Topology file for the hinge-like particle.

