## Supplemental figures for "Particle deformability enables control of interactions between membrane-anchored nanoparticles"

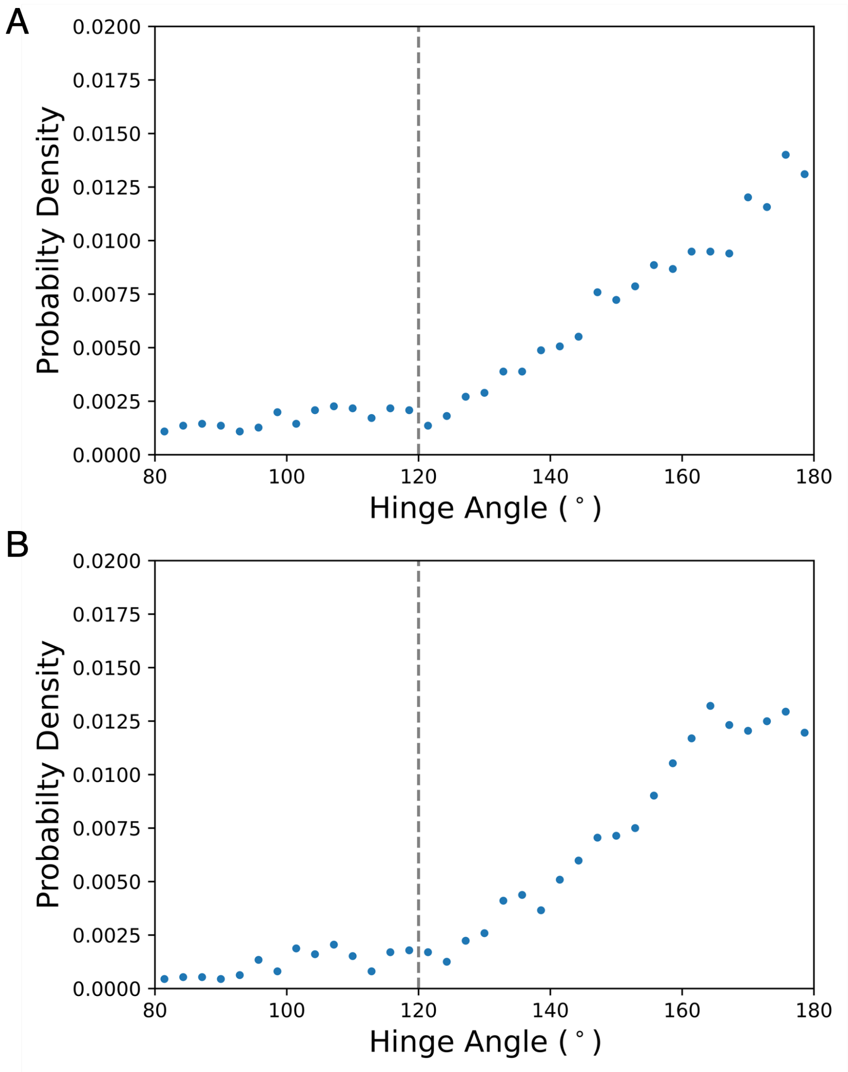


Figure 1: The probability density of the hinge angle ($\theta$) at equilibrium for a hinge in water with $k = 0$. (A) With cholesterol anchors. (B) Without cholesterol anchors.

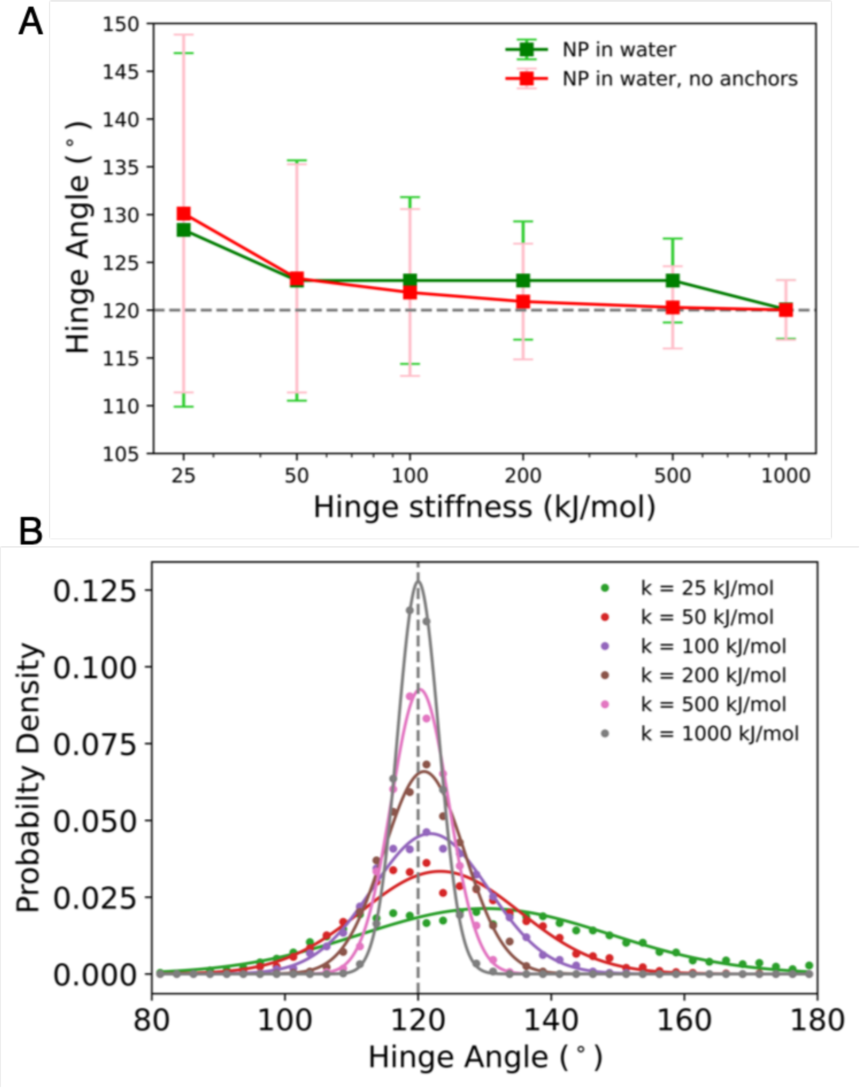


Figure 2: (A) Comparison of the angles sampled by isolated hinges in water with and without cholesterol anchors. The horizontal dashed line corresponds to the natural angle of the hinge ($\theta_0$). (B) The probability density of the hinge angle ($\theta$) at equilibrium for a hinge in water without cholesterol anchors. Different values of the hinge stiffness ($k$) are shown. Solid lines are Gaussian fits to the data (circles) meant to guide the eye.

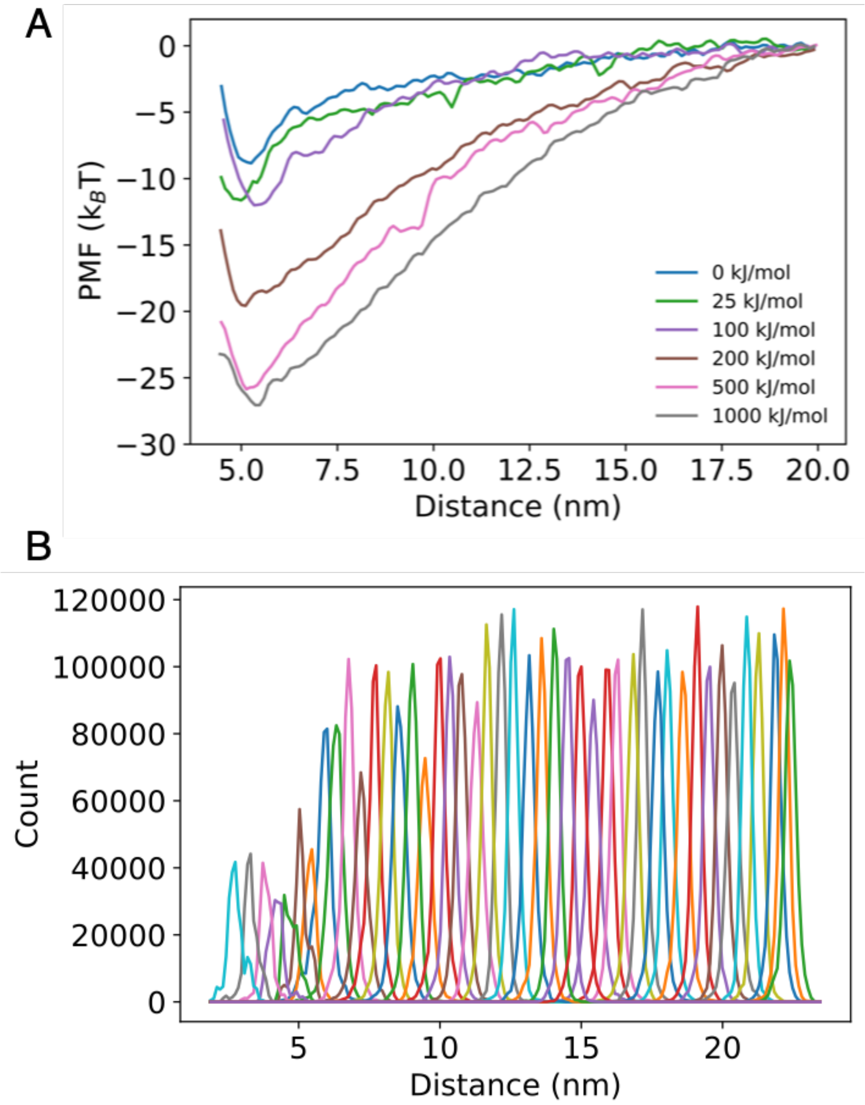


Figure 3: (A) Potential of mean force as a function of the distance between the centers of mass of the two hinge-like particles in the plane of the membrane ($xy$-plane). (B) Histograms obtained for different windows during umbrella sampling. The case shown is for the stiffest hinge with $k = 1000$ kJ/mol.

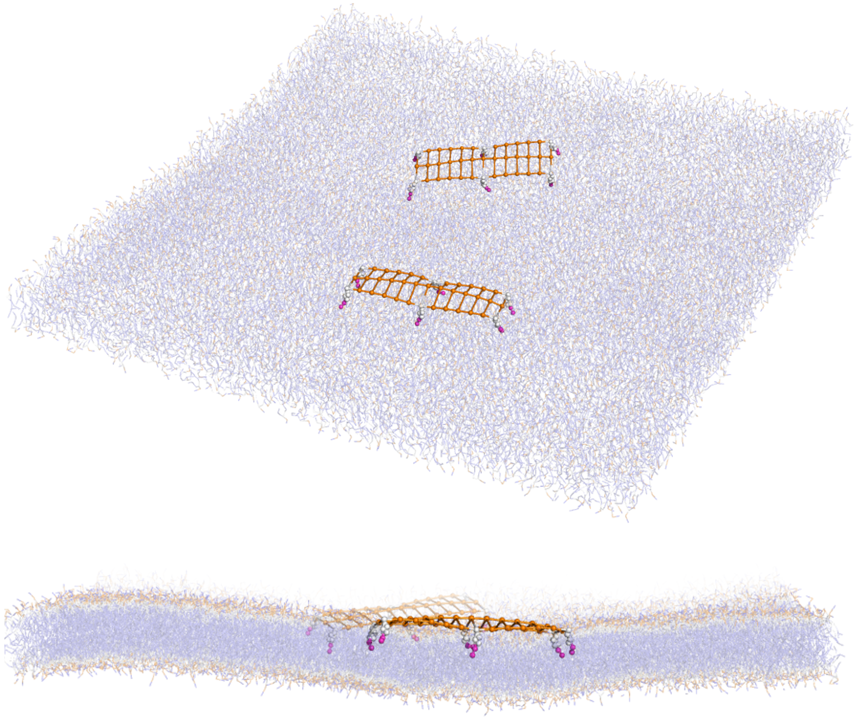

Figure 4: Simulation snapshot of two hinge-like particles anchored to a membrane with $k = 25$ kJ/mol, viewed from different angles. This demonstrates how the particles become slightly distorted when the cholesterol anchors insert into the membrane.
